# Rapid and reusable high-throughput microfluidics through modular assembly

**DOI:** 10.64898/2026.01.12.699088

**Authors:** Linh T.P. Le, Omkar Hegde, Wei-Huan Wu, Ayesha Ejaz, Abhisek Dwivedy, Xing Wang, Minjun Son

**Affiliations:** Biohub Chicago, Chicago, IL, USA; Department of Bioengineering, University of Illinois at Urbana-Champaign, Urbana, IL, USA; Pritzker School of Molecular Engineering, University of Chicago, Chicago, IL, USA; Nick Holonyak Jr. Micro and Nanotechnology Laboratory, University of Illinois at Urbana-Champaign, Urbana, IL, USA; Carl R. Woese Institute for Genomic Biology, University of Illinois at Urbana-Champaign, Urbana, IL, USA; Department of Chemistry, University of Illinois at Urbana-Champaign, Urbana, IL, USA; Cancer Center at Illinois, University of Illinois at Urbana-Champaign, Urbana, IL, USA; VinUni-Illinois Smart Health Center, VinUniversity, Hanoi, Vietnam

## Abstract

High-throughput microfluidics has transformed biomedical research by enabling precise and parallel sample handling, but most devices are single-use due to channel occlusion and contamination from experiments. Alongside low fabrication yield and reduced experimental success associated with dense microfeatures, this creates a major bottleneck for scalable high-throughput applications. We present a rapid, reusable, and modular high-throughput microfluidic platform with integrated microvalves for automation. The platform employs a multilayer architecture consisting of a custom casing, PDMS layers with dense microfeatures for fluid handling and culture, and a glass substrate. Permanent bonding is applied only between control and fluid layers, while reversible bonding is used at all other interfaces, including the substrate. Because substrate is the primary cell-contact surface and can be readily detached, the remaining layers can be disassembled, thoroughly cleaned, and reused with minimal processing on a new substrate. This approach improves repeatability and experimental success while reducing preparation time from days to ∼2 hours. The disassemblable design also supports incorporation of application-specific layers between fluid layer and substrate, enhancing platform versatility for 3D culture. We validated performance through pressure/flow characterization and on-chip cell/organoid culture. Overall, our platform accelerates rapid high-throughput data generation across diverse biological applications.

## Introduction

Microfluidic technologies enable precise handling of fluids and samples at picoliter to micrometer scales, accelerating advances in biological research and medical devices [1, 2]. By integrating functions such as droplet generation, mixing, multiplexing, microvalve, and cell trap into lab-on-chip platforms, microfluidic systems enable experiments that are difficult to achieve using conventional approaches while also supporting high-throughput operation through multiplexing and parallelization [3–5]. For example, droplet-based microfluidics enables the rapid encapsulation of single cells with biomolecules in picoliter reactors or droplets for massive drug screening [6, 7], while microfluidic mixers facilitate rapid and well-controlled chemical reactions [8, 9]. Multi-layer microvalve technologies have been widely explored to precisely regulate chemical compositions or gradients in nanoliter size biological cultures, as well as to increase throughput through multiplexed designs [10–12]. Organ-on-chip platforms further extend these capabilities by recreating physiologically relevant microenvironments that mimic tissue-level functions [12–15]. Collectively, these advances provide unprecedented throughput, automation, and spatiotemporal control over experimental conditions, reducing reagent consumption while increasing reproducibility and experimental precision. As a result, microfluidics has become a powerful tool across diverse areas of biological research, including single-cell analysis, drug screening, and systems biology.

To maximize throughput, microfluidic systems rely on small and closely spaced features that are often on the order of a few tens of microns, or even submicron in scale, resulting in high integration density [7,8]. Designing and operating chips at this scale enables a high degree of parallelization, as many independent channels, chambers, or reaction units can be replicated and operated simultaneously. Large arrays of microchambers or parallel flow channels allow concurrent biochemical assays, cell culture experiments, or single-cell analyses to be performed on a single chip [6, 7]. In addition to this spatial parallelization, droplet microfluidic platforms achieve high-throughput through rapid temporal parallelization, enabled by small, closely spaced channel geometries that support fast fluid switching and stable droplet breakup at high frequencies; as a result, millions of droplets can be generated sequentially [12, 13, 16]. More broadly, the micro-sized features in microfluidics are well matched to the dimensions of biological cells, enabling precise manipulation, confinement, and interrogation of individual cells or small cell populations [17, 18]. Channel and chamber dimensions comparable to cell size allow controlled microenvironments, efficient chemical deliveries, and single-cell–resolved measurements that are difficult to achieve in macroscale systems [10–12, 15]. Collectively, the high integration density of microfluidic systems facilitates throughput and precision in biological studies.

Automation is another key technology employed to leverage the high-throughput capability of microfluidic systems. Due to a large number of samples each requiring rapid and precise control of fluid flow, manual operation becomes impractical, particularly when dynamic and continuous modulation of individual sample condition is desired [19–21]. One commonly used approach to achieve this level of control is the two-layer microvalve scheme, in which a control layer containing pneumatic channels is separated from a flow layer by a thin deformable membrane, typically made of Polydimethylsiloxane (PDMS) [10, 11]. By pressurizing the control channels, the membrane deflects to locally open or close the underlying flow channels. When combined with electronically addressable solenoid valves that actuate each microvalve according to user-defined programs, this approach enables complex yet precise regulation of fluid routing, timing, and volume across hundreds of samples with minimal error [10, 11, 22].

Although microscopic and dense features, precise control by multilayer microvalves, and automation through programmable solenoid valves offer substantial benefits, the integration of these elements also increases system complexity and the likelihood of failure during both chip fabrication and operation [22–27]. Producing molds for devices with high feature density requires advanced micro-/nano-fabrication techniques to ensure high pattern fidelity and low defect rates [25–27]. Additional challenges arise when fabricating multilayer devices from these molds, where densely packed fluid channels must be precisely aligned with a large number of smaller and more densely arranged control lines for microvalve operation [27–29]. As feature density increases, device performance becomes less tolerant of microscopic defects: even small dust particles can cause channel blockage or crosstalk, and subtle PDMS deformation can lead to partial misalignment, compromising overall chip functionality. Microfeatures further exacerbate experimental failure during device operation. Narrow microchannels inherently increase hydraulic resistance, necessitating higher pressures to achieve target flow rate [30–32]. Furthermore, microchannels with small cross-sectional areas are highly susceptible to clogging by debris or contaminants introduced with cell suspensions or other perfusion media. Consequently, microfluidic devices with dense microfeatures often exhibit low fabrication yield, and experimental success rates remain limited even for otherwise functional chips.

Together, fabrication challenges and operational failures result in an inefficient workflow that undermines the theoretical advantages of high-throughput microfluidics. Moreover, despite the substantial time, effort, and resources required to achieve a successful experiment, many devices are effectively single-use. Once a chip becomes clogged or contaminated, cleaning and reusing are difficult due to the confined and microscale nature of the microfluidic channels. Additionally, many devices are designed and built for specific applications. Even minor changes in the experimental objectives often require redesign and refabrication of the molds and chips. Collectively, these limitations hinder the routine adoption of high-throughput microfluidics and largely confine such platforms to specialized and well-funded engineering laboratories.

To overcome these limitations, we introduce a modular design strategy that decouples the sophisticated control and fluid infrastructure from the easily contaminated sample substrate (Fig 1A, B). Our modular design enables direct and thorough cleaning of fluid channels, allows clean reuse of the platform within a few hours, and facilitates easy modification of the substrate to accommodate diverse biological applications (Fig 1C). Importantly, we demonstrate that these advantages are achieved while preserving robust microvalve functionality and maintaining highly dense microfeatures to maximize throughput, thereby facilitating broader and robust adoption of microfluidics for high-throughput data generation.

**Fig 1.**
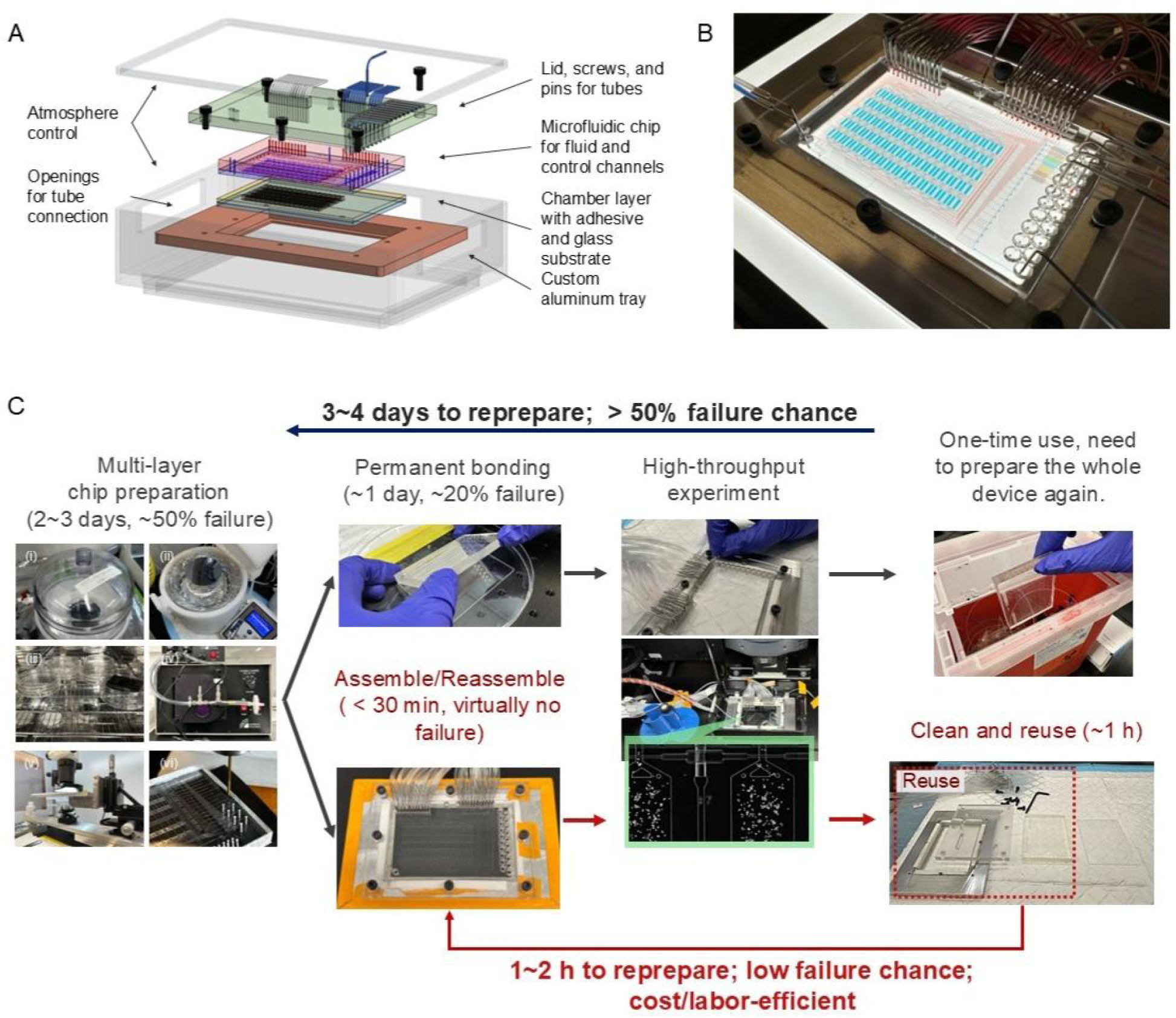
Design and workflow of the modular microfluidic platform. **A.** Schematic of the device and platform assembly illustrating the multilayer components. The complete assembly is designed to fit within an atmospheric control box for cell culture. **B**. Picture of a fully assembled device loaded with food dye. Red indicates control channels, while blue indicates fluid channels and cell chambers. **C.** Flowchart comparing the proposed modular assembly method (red arrows) with the conventional approach (black arrows). For multilayer chip fabrication, mandatory steps include (i) mixing/degassing PDMS for control layer, (ii) PDMS spin coating for fluid layer, (iii) overnight baking, (iv) surface plasma treatment; (v) aligning and bonding; (vi) hole punching. Our modular design mitigates assembly-related failures and enables rapid reuse of the chip within 2 h, thereby substantially increasing experimental success rates and effective throughput.

## Results

### Modularization facilitates rapid and efficient microfluidic experiments while enhancing platform versatility

While developing a reusable high-throughput platform, one key challenge was how to maintain the bonding between layers during experiments where high pneumatic pressure is required [10]. Especially as chip designs become more complex and channels become smaller and more densely packed, the pneumatic pressure required to achieve the necessary flow rate increases, while the relative bonded surface area available to withstand pressure in the flow channels decreases.

To address the tradeoff between channel density or experimental throughput and the surface area for bonding we modularized the platform, decoupling the reusable fluid-control infrastructure from the disposable sample environment (Fig. 1A and B). At the core, the system consists of two modules: a fluid-control module and a replaceable substrate. The fluid-control module is further divided into two microchannel layers separated by a thin (∼20 µm) PDMS membrane – control channels on the top and fluid channels on the bottom – to route specific input media to designated locations or cell chambers [10]. Channels in both layers incorporate a multiplexing architecture, enabling operation of many pathways with a minimal number of control lines. Because two-layer microvalves often require more than 10 psi of pneumatic pressure to fully close the underlying fluid channel, robust bonding between the two layers is essential [10]. We therefore applied plasma bonding to ensure strong and reliable adhesion between the control and fluid layers (Fig. 2A) [10]. Although this confines the control channels to be in closed geometry with dead-ends, only deionized (DI) water is loaded into the control channels, and we confirmed the entire fluid-control module is autoclavable, mitigating contamination concerns. When assembled, the fluid-control module sits on a replaceable substrate, clamped between a transparent polycarbonate (PC) lid and a custom-designed aluminum tray that fits inside the incubation box for culture atmosphere control (Fig. 1A).

**Fig 2.**
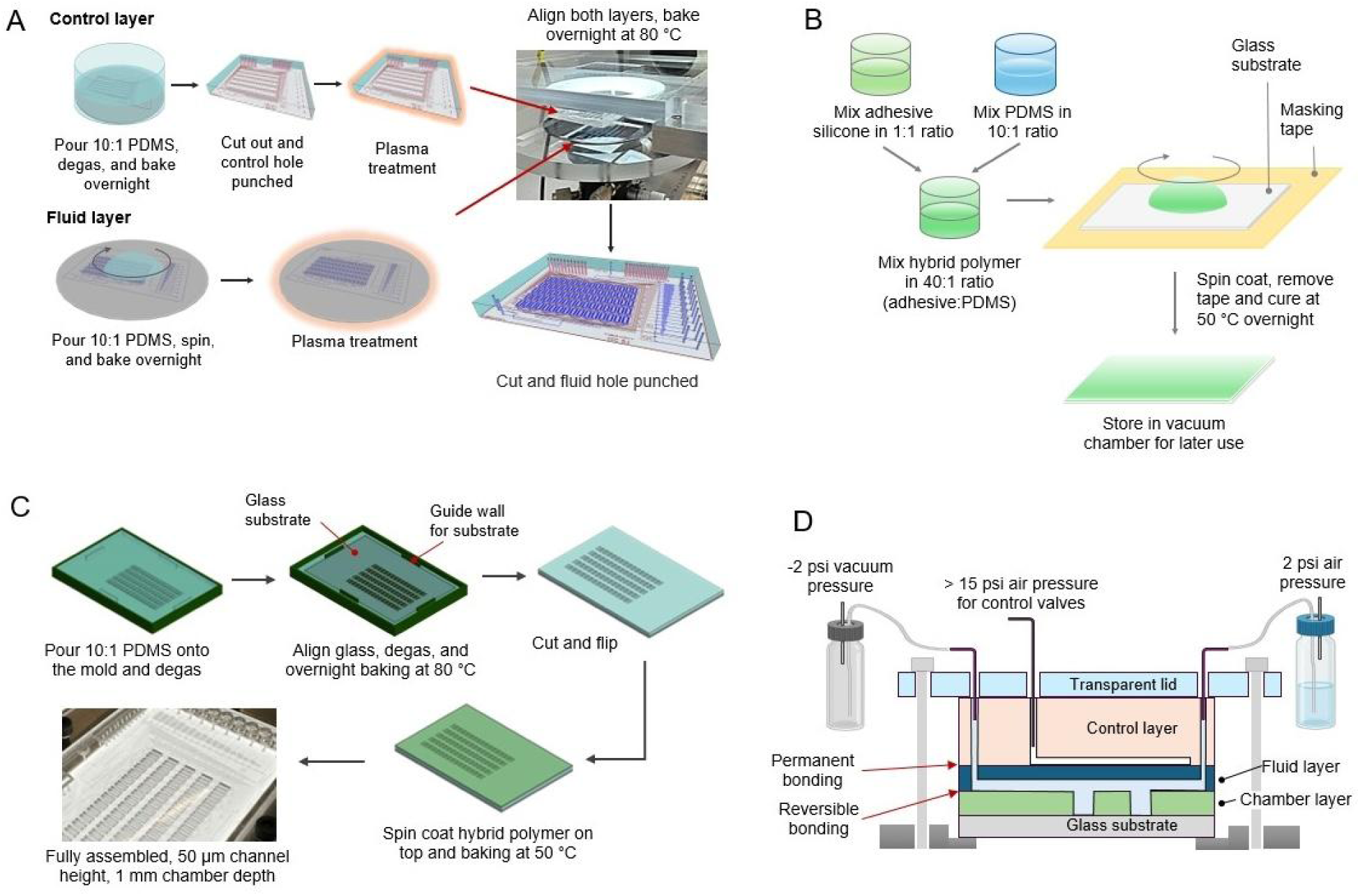
Fabrication and assembly of the modular microfluidic platform. **A.** Schematic workflow for fabricating the reusable PDMS fluid-control module. Control holes are punched prior to alignment. **B.** Workflow for the thin-substrate. A hybrid adhesive polymer (mixture of adhesive silicone and PDMS) is spin-coated onto a glass slide and cured to create a reversible sealing interface. **C.** Workflow for the deep-well substrate for 3D culture. Using a 3D-printed master, 1 mm deep chambers are molded in PDMS with a glass slide on top. **D.** Cross-sectional diagrams of the final device assembly. The fluid-control module and deep-well substrate are manually aligned using outlines incorporated into the design and the corresponding inset in the aluminum tray, then mechanically clamped. Negative pressure can be applied to further increase flow rate.

The reusable fluid-control module is fabricated using a workflow that is slightly modified from previous designs (Fig. 2A) [10, 11]. Briefly, the control-layer – which comprises most of the chip body – is cast from 8:1 (base:curing agent, by weight) PDMS. The layer is trimmed to match the substrate outline, and access ports for the control lines are punched. The control layer is then oxygen-plasma treated, aligned, and bonded to the fluid layer, which consists of 15:1 PDMS spin-coated on the fluid-channel mold (∼ 50 μm thickness). After bonding, the assembled chip is trimmed again to match the substrate, and the remaining access ports to the fluid channels are punched. In addition to using different PDMS mixing ratios to tune layer stiffness for control layer and promote bonding between the fluid layer and substrate, a key difference from prior workflows is that the holes to access control lines have to be punched prior to alignment to the PDMS spin-coated fluid mod. This procedure ensures there is ∼ 50 μm PDMS membrane permanently bonded beneath each control hole to reduce the impact of high pneumatic control pressure on the more weakly bonded substrate.

Although simply mounting this fluid-control module on a clean non-treated glass substrate with mechanical clamp can work at low fluid-layer pressure (< 2 psi), we found that adding a surface coating to the substrate improves robustness and enables higher fluid-layer pressures to achieve the increased flow rates needed for faster chip operation (Fig. 2B) [33, 34]. Among the coating methods we tested, a hybrid adhesive polymer performed best, providing robust reversible bonding while maintaining a clean, smooth surface for microscopy and a sterile environment for cell culture. More details are discussed in the next section. However, regardless of substrate composition, full device operation requires mechanical clamping via the polycarbonate lid and aluminum tray. When high pneumatic pressure (>15 psi) is applied to the control lines to close the fluid channels, it not only deforms the fluid layer but also increases the interfacial stress between the fluid layer and substrate, promoting delamination. In practice, we often observed delamination around the control-line access holes, even with mechanical clamping and the adhesive polymer. A key design challenge, therefore, was minimizing how this delamination propagates to or disrupts the fluid channels, which we discuss in the following sections.

Beyond spin-coated glass substrate, our modular design also supports interchangeable substrate formats that interface with the same fluid-control module, depending on the experimental requirements (Fig. 2C and D). For example, the spin-coated glass substrate described above is more suitable for high-resolution 2D imaging for single-cell experiments, whereas a deep-well PDMS substrate molded from a 3D printed master provides more suitable platform for applications such as 3D organoid culture in gel or for reducing fluid-induced shear stress on cells within the chamber. The deep-well substrate also enables gel or cell loading prior to device assembly, thereby avoiding delivery of cells or viscous media through microchannels. Channel congestion during cell loading has been a recurring limitation of previous high-throughput microfluidic chips. By loading cells from outside the device, analogous to seeding a well plate, this approach helps keep the channels clean and ensure the channels function as designed throughout the experiment.

Overall, unlike conventional monolithic devices in which all layers are permanently bonded and the chip is discarded after a single use, our approach enables repeated disassembly and reuse of the fluid-control module without refabricating the device. Moreover, through modularization, the substrate can be readily customized to accommodate diverse culture requirements. In the following sections, we demonstrate that these innovations are compatible with complex high-throughput microfluidic designs featuring dense multilayer networks of microscopic channels operated across a wide range of pneumatic pressure.

Our workflow analysis highlights the efficiency gains of this modular strategy (Fig. 1C). Conventional multilayer fabrication typically requires a 3–4 day cycle and accumulates failure risk across mold preparation, PDMS casting, layer alignment, and plasma bondings. In contrast, once the fluid-control module is validated in our approach, subsequent experiments require only a new substrate, which is inexpensive and can be mass-produced and stored. Replacing the substrate from a prepared batch reduces per-experiment setup time to <2 hours and lowers the likelihood of failure, since a substrate can be readily swapped or realigned on site if needed. Hence, our method substantially reduces the preparation time, improves reliability, and enhances versatility by allowing reuse of the most failure-prone and labor-intensive component.

### Adhesive polymer enhances necessary reversible bonding for stable chip operation

Reliable and robust operation is essential for high-throughput microfluidic devices, which must control fluids across multiple channels in a precise and consistent manner over extended periods. A key challenge lies in achieving a robust yet reversible seal at the fluid layer and substrate interface under elevated fluid pressures. This is particularly difficult in high-throughput designs, where channel densities are high and adjacent channels may be separated by gaps as small as ∼100 μm, limiting the available bonding area to withstand the pressures applied within the channels. In our design, increasing fluid pressure revealed that leakage typically initiated at regions where higher local pressure is anticipated, for example, at junctions between several channels where microvalves are concentrated or near access holes for inlet fluids (Fig. 3A).

**Fig 3.**
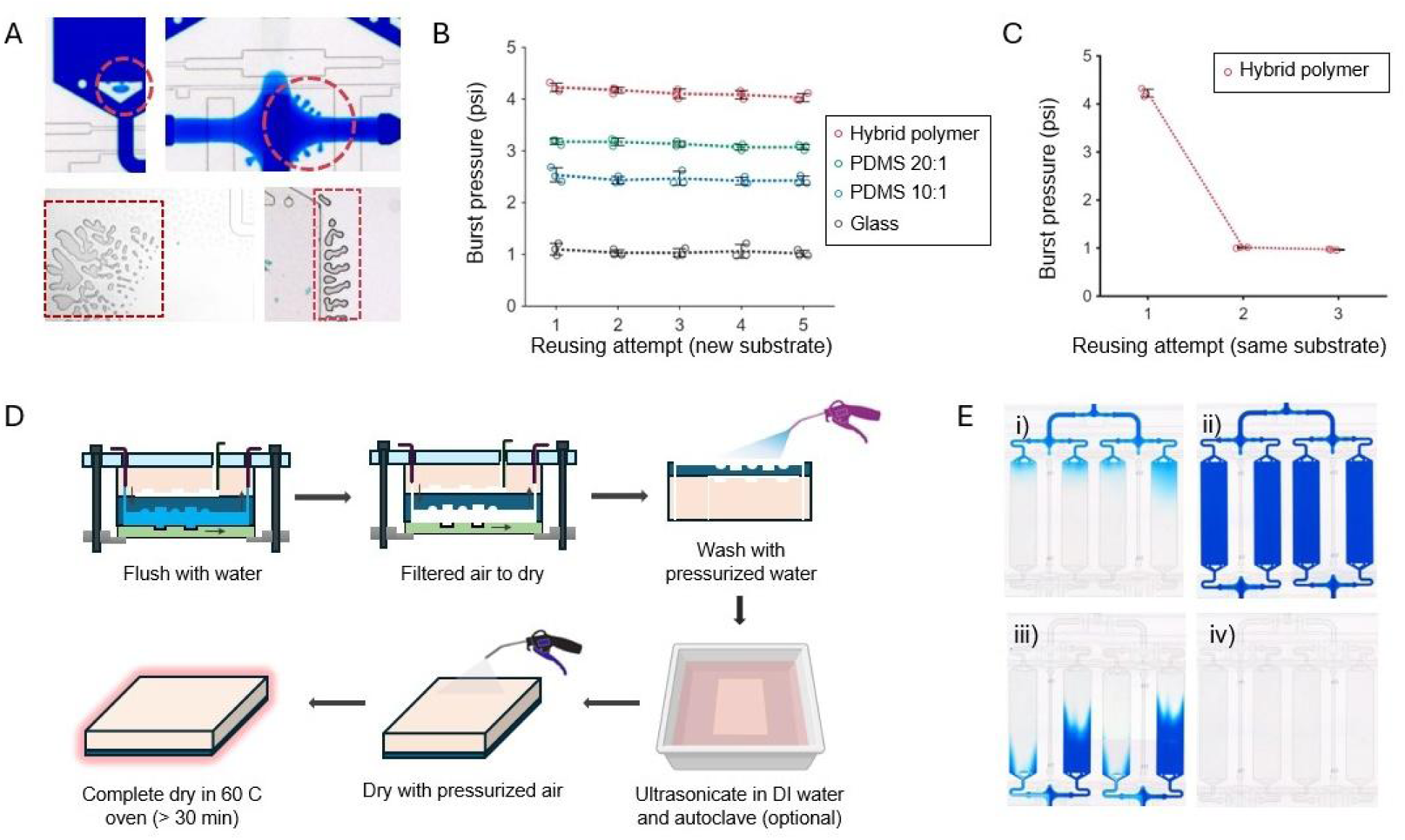
Characterization of reversible bonding and the cleaning workflow for device reuse. **A**. Images showing representative delamination at the substrate interface (red dotted circles and rectangles) observed during burst testing. **B.** Error bar plots showing burst pressure measurements for glass substrates spin-coated with hybrid adhesive polymer, 10:1 PDMS, 20:1 PDMS and uncoated (plain) glass. For each reuse, the substrate was replaced while the same PDMS body was reused after washing. Circles denote the three replicate trials for each substrate at each reuse, while the dotted line connects the mean of the three trials across reuse. **C**. Similarly, burst pressure was measured for glass substrates coated with the hybrid adhesive polymer; however, this time the same substrate from the prior run was reused rather than replaced with a newly coated one. **D.** Illustration of the washing and re-preparation protocol for the fluid-control module. Ultrasonication and autoclaving are optional. **E.** Series of microscope images with blue dye to demonstrate the effectiveness of Teflon surface coating. Channels filled with blue dye (i, ii) were successfully flushed, leaving a residue-free surface (iii, iv).

To achieve a reliable and reversible bonding at the substrate interface, we evaluated a few different surface-coating methods using our platform (Fig. 3B). Briefly, glass substrates were spin-coated with either PDMS (10:1 or 20:1 base:curing agent, by weight) or a hybrid adhesive polymer consisting of 40:1 mixture of 1:1 adhesive silicon and 10:1 PDMS. In addition to these, an untreated glass slide was included as a negative control. For delamination testing, we first closed the microvalves for all inputs and outputs, while keeping all other microvalves – including those in the multiplexer, chamber controls, and bypasses – open. We then pressurized a single input to the target pressure, opened the corresponding valve, waited several minutes, and inspected the device for delamination. Because all flow paths were open and terminated in dead ends under this configuration, the applied pressure was expected to equilibrate throughout the device, enabling assessment of delamination anywhere on the chip. Among the substrates tested, the hybrid adhesive polymer performed best, withstanding up to 4.2 – 4.5 psi. The 20:1 PDMS coating also performed well, tolerating pressures up to 3.2 psi, which is sufficient for most applications especially with negative pressure applied on outlet (Fig. 2D).

Next, we evaluated reusability of the fluid-control module by replacing the substrate with a newly fabricated one for each subsequent run (Fig. 3B). The burst pressure remained consistent across repeated trials, indicating that performance is maintained over multiple uses and confirming the reusability of the platform. In contrast, reusing the same adhesive-coated substrate from a prior run led to a significant reduction in burst pressure to ∼1 psi (Fig. 3C), likely due to deformation or tearing of the compliant adhesive layer during disassembly (Fig. S1). This result is consistent with prior reports of this material’s mechanical properties [33, 34]. Consequently, we prioritized reuse of the fluid-control module while treating the substrate as a low-cost and disposable component that can be fabricated in batch and readily replaced as needed.

Additionally, to minimize the cross-contamination from repeated use, microchannels in the fluid layer were coated with Pluronic 0.2% or Teflon 0.5% prior to experiments to reduce nonspecific adsorption [10, 35]. After each run, channels were flushed with DI water followed by air, and the device was then disassembled, leaving the bottom surface of the fluid channel exposed to air (Fig. 3D). This provided direct access to the fluid channels for thorough cleaning with pressurized DI water. After inspection with a microscope, if additional cleaning was deemed necessary, we applied ultrasonication and autoclaving to ensure the channels were clean and sterile. Following rinsing, the module was dried with filtered pressurized air and then incubated at 60°C for more than 30 min. This protocol effectively removed residual reagents and restored the device to its original performance, as confirmed by dye-based visualization (Fig. 3E). The fluid-control module was then ready for reassembly with a fresh substrate, enabling rapid turnaround between experiments.

### Chip designs isolating higher-pressure pathways improve platform reliability during complex and extended operation

In high-density multilayer microfluidic systems, strong plasma bonding is typically necessary for maintaining structural integrity under elevated operating pressures. Although we characterized the burst pressure of our modularized platform and confirmed that the chip does not delaminate under operational fluid and control pressures (Fig 3B), it remained unclear whether the platform would continuously perform reliably and as designed during long and complex high-throughput experiments. During these experiments, microvalves repeatedly actuate, and chambers and flow paths undergo frequent pressurization and depressurization.

In addition to the adhesive polymer coating and mechanical clamping described above, we strategically optimized the device layout to improve operational robustness (Fig. 4A). Briefly, we increased the distance between the fluid inlet holes and the high-pressure regions of the device to provide sufficient bonding area and to minimize deformation from tubing inserted into the inlets. Additionally, we clustered the access holes for the control lines and spatially separated them as far as possible from the fluidic components. Although this layout led to more pronounced local delamination around the control-hole cluster, where high-pressure control lines are concentrated, the thin PDMS layer permanently bonded underneath mitigated its impact, and the clamping system prevented propagation of delamination beyond the vicinity of the cluster. Together, these design principles substantially improved overall platform robustness. Other newly implemented features include serpentine channels around the fluid inlets to equalize hydraulic resistance and flow rates across inlet media (Fig. 4A-i), as well as multiplexing implemented along both layout axes to further increase the number of samples (Fig. 4A-ii). For comparison, although the overall chip size is similar to our previous conventional chip designs, the new design increases chamber area by ∼2 fold (1.8 vs 3.4 mm^2^), allowing more cells to be tested per sample, while doubling the number of chambers (64 or 92 vs 160 individual controllable chambers) for higher throughput [10, 36–38]. Together these design principles ensure that the sophisticated fluid-control infrastructure remains functional in our modular platform while enhancing through-put.

**Fig 4.**
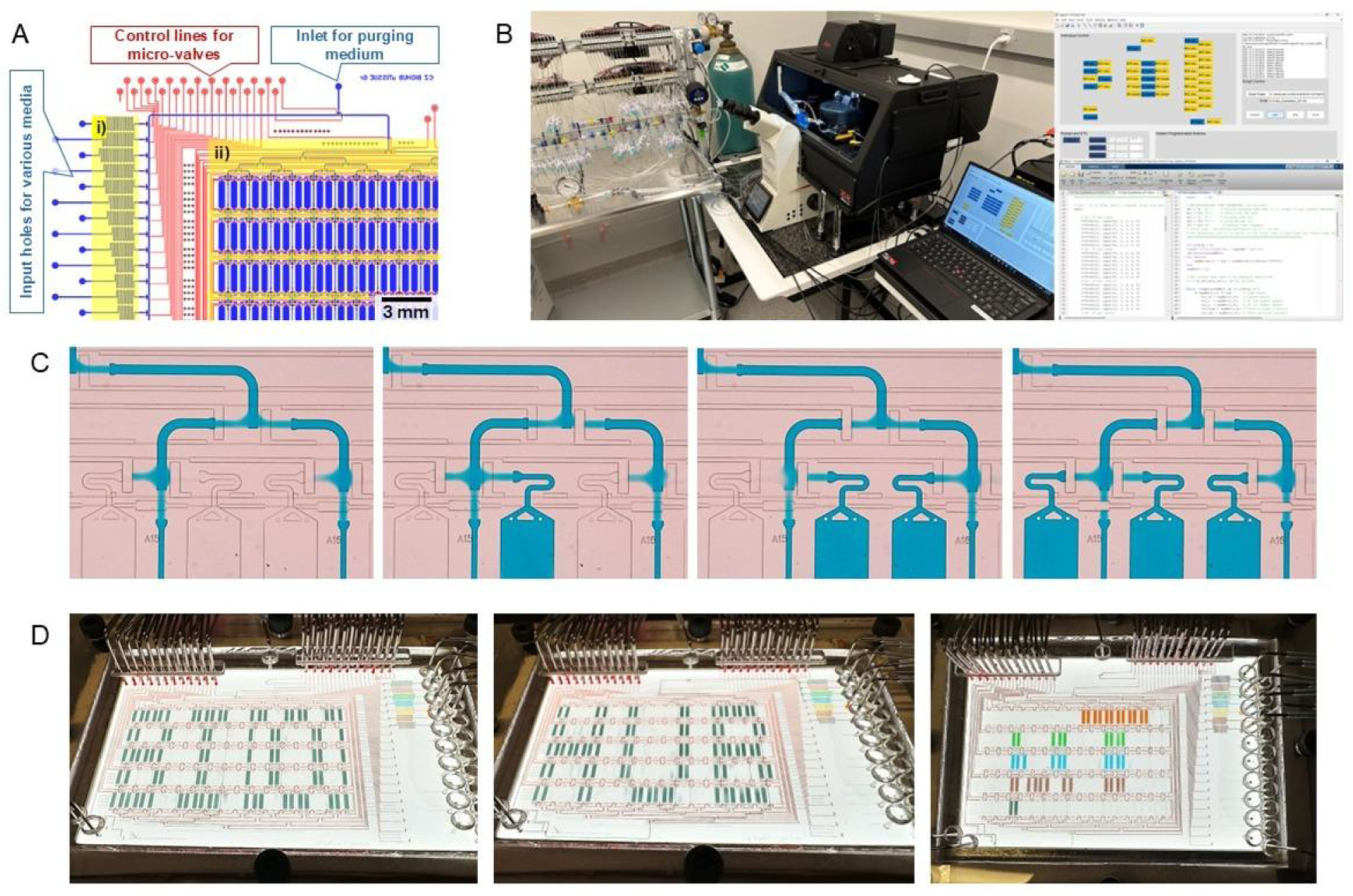
Core chip design, automation, and functionality test. **A**. CAD design of the fluid (blue) and control (red) channels, including serpentine paths to match fluid resistance from each inlet (i), multiplexers (ii). **B**. Automation and imaging setup, including pneumatic solenoid valve manifolds to actuate control lines, an inverted microscope with motorized stages, and custom software to operate solenoid valves according to prewritten schedule. **C**. Time-lapse images showing fluid handling on our platform and robust microvalve operation to sequentially close and open designated flow paths, directing medium (blue dye) to specific chambers without affecting neighboring chambers. **D**. To validate the reliability of our reusable, modular high-throughput platform across the entire chip, we selectively and sequentially filled each chamber with colored dyes to create distinct patterns, and observed no cross-contamination.

To validate the automation fidelity and long-term reliability of the integrated system, we developed custom software with a graphical user interface (GUI) (Fig. 4B) and performed extended routing operations (Fig. 4C,D). The system reliably executed automated fluid-handling sequences, including multiplexer reconfiguration, bypass feeding, simultaneous chamber opening/closing, and sequential single-chamber feeding. Across all operations, microvalves repeatedly actuated to open and close the underlying flow channels without any noticeable failures (Fig. 4C).

To further assess fluid-handling precision and long-term operational robustness, we ran a programmed feeding schedule that generated a series of spatial patterns across the chamber array (Fig. 4D). The platform consistently addressed individual culture units, selectively loading colored dyes to form distinct letter-shaped patterns. We observed no cross-contamination between adjacent chambers, supporting the platform’s suitability for high-throughput biological workflows.

### Single-cell experiments demonstrate the potential of the reusable microfluidic platform for continuous high-throughput data acquisition

A key advantage of a chamber-based high-throughput platform is the ability to precisely deliver a specific medium or stimulus to each chamber and to completely exchange medium in each chamber with minimal perturbation to cells, while enabling high-quality imaging with low background fluorescence, owing to the shallow and uniform chamber design. Each chamber can be perfused precisely and subsequently according to a user-defined schedule, with media exchanges executed at exact preprogrammed time via custom software.

To demonstrate these capabilities in our modular and reusable platform, we loaded mouse macrophages (RAW 264.7) into chambers in the thin-substrate configuration and sequentially exposed them to a range of lipopolysaccharide (LPS) doses (Fig. 5A). We then monitored NF-κB pathway responses in macrophages under this dynamic LPS stimulation using fluorescence imaging (Fig. 5B-D). NF-κB is a key transcription factor that regulates innate immune responses downstream of cytokines and pathogen-derived cues, and is known for its oscillatory nuclear translocation dynamics (Fig. 5B) [38–40]. Previous studies have shown that features of these oscillations, such as oscillation frequency and changes in peak amplitudes, shape downstream gene regulation and, consequently, immune responses [41–44]. Several studies have further examined how the oscillatory behavior of NF-κB translocation depends on temporal changes in extracellular cues [45, 46]. For example, our previous work tested stepwise changes in TNF-α concentration in 3T3 mouse fibroblasts and found that NF-κB responses track changes in TNF-α rather than the absolute dose at a given time [45]. In contrast, stimulation with Pam3CSK4 (PAM), a synthetic lipopeptide mimicking bacterial lipoproteins, elicited a qualitatively different response [46]. After a single strong activation, NF-κB hardly responded to further increase in PAM concentration, highlighting that NF-κB signaling dynamics can vary substantially across ligands. Our platform is well suited for probing how diverse ligand combinations and temporal stimulation patterns regulate NF-κB dynamics across different cell types.

**Fig 5.**
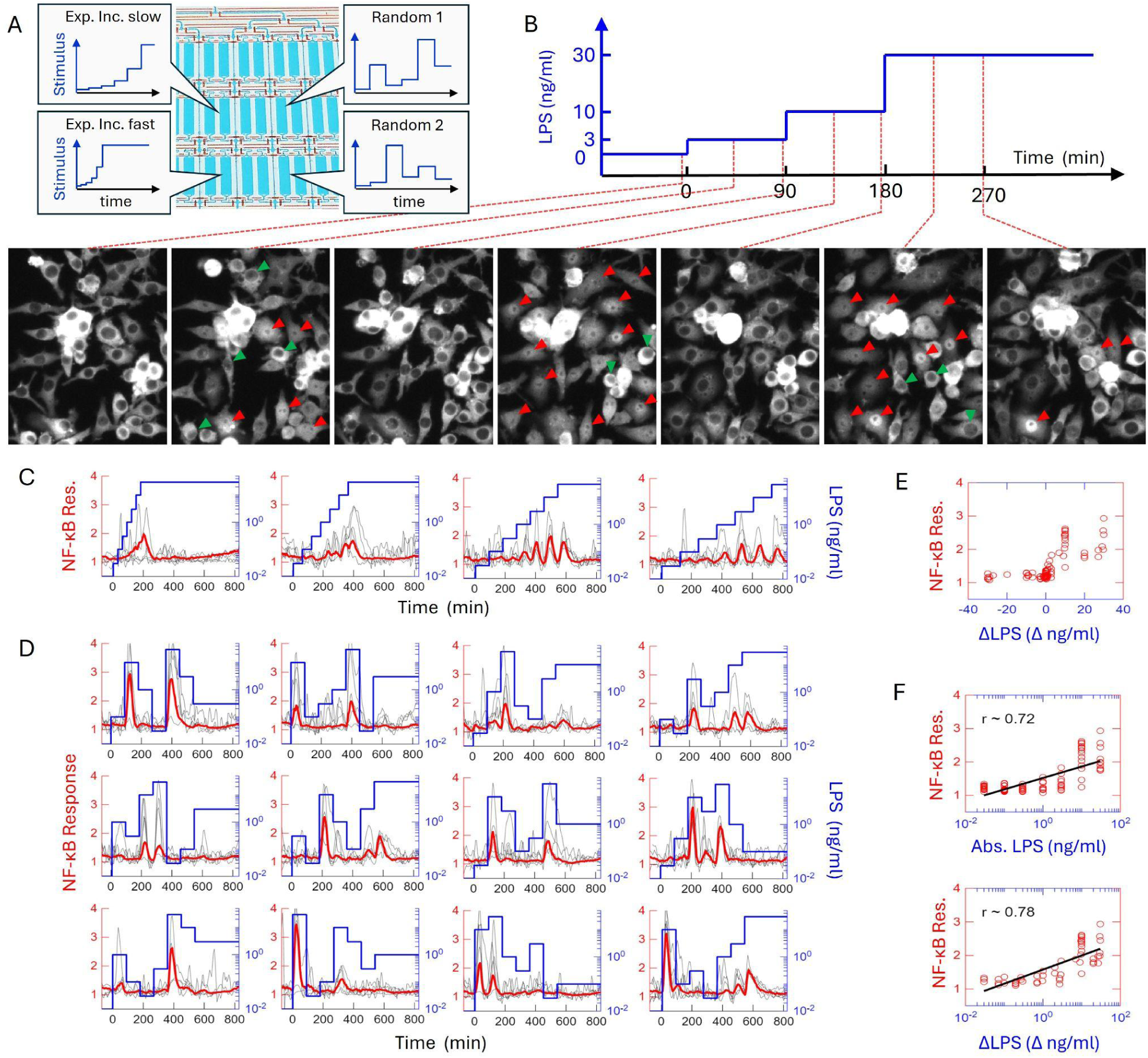
NF-κB responses to dynamic LPS stimulation as a demonstration of our modular platform. **A.** Schematic of the multiplexed experimental design, illustrating four exemplary LPS stimulation profiles (slow and fast exponential increases, and two randomized patterns). **B.** Representative time-lapse fluorescence images of NF-κB reporter macrophages during stepwise LPS increases (0 to 30 ng/mL). Red and green arrowheads indicate examples of activated and non-activated macrophages, respectively, after each increase. **C.** NF-κB activation, quantified as the median nuclear fluorescence of the NF-κB reporter, in response to exponentially increasing LPS at different rates. Thin gray lines show responses from 10 randomly selected single cells; the thick red line shows the population median; the blue line shows the LPS dose in log scale. **D**. Similarly, NF-κB responses were measured under 15 randomly fluctuating LPS dose patterns (more shown in Fig S3). **E.** Scatter plot comparing the subsequent NF-kB response with the change in LPS doses (ΔLPS) across all samples in panel D. **F.** Correlation analysis comparing NF-κB responses against absolute LPS concentration (top, r ∼ 0.72) versus change in LPS concentration (bottom, r ∼ 0.78).

When we applied an exponentially increasing LPS dose to macrophages expressing a fluorescent NF-κB reporter, we observed a response pattern reminiscent of fibroblast NF-κB dynamics under exponentially increasing TNF-α (Fig. 5B,C, and S3). Although this trend was weaker and persisted for a shorter duration (less than 10 h post the initial stimulation), NF-κB activity nonetheless continued to rise or remained elevated as the LPS dose increased exponentially. This behavior is especially notable when compared with fibroblast responses to PAM, another pathogen-derived ligand similar in origin to LPS, where increasing PAM elicits little oscillatory behavior and instead produces a single dominant activation peak [46]. Consistent with this, prior work also reported that NF-κB in 3T3 fibroblasts does not exhibit robust oscillations under high-dose LPS stimulation [45]. In contrast, in our experiments NF-κB in macrophages displayed pronounced oscillatory dynamics under increasing LPS stimulation.

Encouraged by these results, we next tested how random temporal fluctuations in LPS dose shape NF-κB response dynamics, reusing the fluid-control module from the previous trial (Fig. 5D and S3). Briefly, cells in each chamber were exposed to 15 randomly ordered LPS doses (0.03, 0.1, 0.3, 1, 3, 10, and 30 ng/mL) with dose changes every 90 min, while acquiring fluorescence images every 6 min. Similar to fibroblast responses to fluctuating TNF-α [45], macrophages responded primarily to increases in LPS dose, while showing little to no response to decreases, largely independent of the absolute LPS concentration after the drop. To better visualize this pattern, we quantified NF-κB peak heights for each interval across all samples and plotted them against the preceding dose change associated with each peak (Fig. 5E). As expected, negative dose changes (ΔLPS < 0) elicited little to no NF-κB response, whereas most positive changes resulted in strong NF-κB activation. To further assess whether NF-κB responses correlate more strongly with the absolute LPS dose or with dose changes, we isolated NF-κB peak heights following only positive LPS changes and plotted them against either the absolute LPS concentration or the magnitude of the dose change (Fig. 5F). We then computed the correlation coefficient for each relationship to evaluate whether NF-κB activity is better explained by absolute LPS dose or by changes in dose. Our analysis revealed that NF-κB response amplitude correlated more strongly with changes in stimulus (ΔLPS) than with absolute concentration, highlighting the importance of stimulus dynamics in immune responses.

Together, these trials demonstrate that our modular platform supports both high-throughput microfluidics and demanding long-term biological experiments, enabling systematic study of diverse conditions and facilitating discovery of new biological phenomena. Despite repeated pressure fluctuations during automated media exchange, the device remained leak-free and mechanically stable. Moreover, cells maintained healthy morphology throughout a 12-hour perfusion experiment, supporting the biocompatibility of the adhesive substrate interface.

## Discussion

Our modular microfluidic platform addresses a key challenge in high-throughput microfluidic research: as device complexity increases to meet the demands of sophisticated biological experiments, reliability, accessibility, and experimental throughput often decrease. In conventional multilayer devices, complex layers are permanently bonded, which concentrates failure risk into a single non-recoverable assembly, typically limits devices to single use, and imposes long fabrication turnaround between experiments [19]. By modularizing these components, our platform enables a reusable, instrument-like fluid-control module that retains the full functionality of multilayer microvalves while eliminating the need to repeatedly fabricate the most complex elements of the device.

The primary advantage of our modular strategy is the significant reduction in experimental cycle time and the significant enhancement of platform versatility (Fig. 1, 2). In conventional workflows, a single failure during mold fabrication, alignment, or irreversible plasma bonding can invalidate an entire 3–4 day fabrication process [19]. In contrast, once validated, the fluid-control module in our system can be reused across numerous experiments, effectively shifting the fabrication effort to the disposable substrate alone. This shift reduces the preparation time for new experiments from days to less than two hours and substantially improves overall reliability. Furthermore, modularity also enables rapid adaptation to different experimental contexts. As demonstrated with deep-well substrates for organoid culture (Fig. 2), the same fluid-control module can be repurposed for distinct biological assays by simply swapping the disposable substrate design.

However, several limitations still remain. Although the fluid-control module is reusable and can be distributed for broader applications, the current setup still relies on custom-built external solenoid valves [10,19], which may be challenging to adopt in standard biology labs. Additionally, although vacuum-assisted outlet operation substantially improves flow rate and stability, some applications benefit from higher inlet pressures (e.g., channel degassing). In our modular system, the maximum operating pressure remains lower than that of permanently plasma-bonded devices (typically >15 psi), which can slow feeding cycles and is not ideal for applications that require high fluid channel pressure [29,33]. Future improvements could incorporate on-chip peristaltic pumping for more precise flow rate control [20] and stiffer backing layers to further increase pressure tolerance. Moreover, while we showed our cleaning protocol and Teflon/Pluronic coating method were effective for soluble reagents [35], experiments involving highly adhesive hydrophobic compounds or sticky cell types may require more rigorous cleaning protocols to prevent contamination between runs.

Despite these drawbacks, we demonstrated that the platform is suitable for its intended biological applications. Specifically, we performed long-term live-cell experiments measuring NF-κB signaling dynamics in macrophages under exponentially increasing and fluctuating LPS stimulation (Fig. 5). In the course of these quick and straightforward demonstrations enabled by our reusable and automated microfluidic system, we uncovered new observations that macrophage NF-κB exhibited robust oscillatory dynamics during LPS fluctuations, responded preferentially to increases in LPS dose, and correlated more strongly with changes in LPS dose than with the absolute dose. In addition, single-cell analysis revealed pronounced cell-to-cell heterogeneity in NF-κB oscillatory behavior under fluctuating LPS stimulation (Fig. 5B–D), consistent with previous studies of NF-κB signaling [40–46]. Notably, cells maintained viability throughout the experiments, indicating that the hybrid adhesive polymer interface is biocompatible and suitable for sensitive cell types [47, 48].

In summary, we developed a reusable and modular microfluidic platform that bridges the gap between fabrication complexity and experimental utility. By integrating a modular architecture with a robust hybrid adhesive interface, vacuum-assisted stabilization, a custom casing that applies uniform sealing pressure, a segregated layout design, and automated control, our plug-and-play approach enables rapid, reliable, and versatile microfluidic experimentation. This platform would provide a scalable framework for diverse applications, including single-cell signaling analysis, organoid culture, and high-content drug screening, while substantially reducing fabrication burden and mitigating failure risk.

## Materials and methods

### Cell culture

Mouse macrophage reporter cells (RAW 264.7 expressing p65-GFP) were cultured in Fluorobrite Dulbecco’s Modified Eagle’s Medium (DMEM; Gibco, A1896701) supplemented with 10% heat-inactivated Fetal Bovine Serum (Omega Scientific, FB11) and 1× GlutaMAX Supplement (Gibco, 35050061) in T75 non-binding flasks. Cells were passaged by washing with phosphate-buffered saline (PBS, pH 7.4; Gibco, 10010023) followed by gentle detachment using a cell scraper. For maintenance, cells were resuspended in fresh medium and split at a 1:10 ratio. Prior to microfluidic experiments, cell suspensions were stained with NucBlue Live (Invitrogen) at a concentration of 2 drops per mL and gently vortexed.

### Microfluidic device design

All microfluidic device components were designed using Autodesk 2025, Onshape, AutoCAD (Autodesk Inc.), or KLayout. 2D designs and 3D print files (.stl) are available in the Supplementary Information.

### Microfluidic experiment setup

Control layer valves were actuated using pneumatic solenoid valves (Festo, 197334) connected via metal pins and water-filled Tygon tubing (ND-100-80) to prevent gas diffusion. Valve timing was automated using a custom MATLAB interface. Media and reagents were delivered to the chip from pressure-relief vials via PEEK tubing (VICI, TPK.505). The microfluidic device was maintained at 37°C, 5% CO₂, and 95% humidity using a stage-top incubator (OKO Lab) mounted on a Leica Thunder microscope enclosed in a custom dark box.

### Control and fluid (reusable) layer fabrication

Master molds were fabricated on silicon wafers following previously described protocols [10].

Polydimethylsiloxane (PDMS) casting was performed using RTV-615 (Momentive) with a base-to-catalyst ratio of 10:1. To fabricate the thick control layer, 60 g of the PDMS mixture was poured over the control mold. For the thin fluid layer, 5–10 g of the mixture was poured onto the fluid mold and spin-coated at 2500 rpm for 1 min. Both layers were cured overnight at 80°C. Following curing, holes were punched in the control layer, and the surface was cleaned using compressed air and adhesive tape. The control and fluid layers were treated with oxygen plasma (Harrick, PDC-001) for 45 s, aligned using a custom stage on an upright microscope (Thorlabs), and bonded. After an additional overnight bake, fluid inlets were punched and cleaned to remove PDMS debris. The assembled chip was bonded to a clean glass slide via oxygen plasma treatment (60 s).

### Fabrication of replaceable layers and chambers

To facilitate replaceability, a hybrid polymer mixture was created by combining a soft silicone adhesive (Liveo™ MG 7-9800 Soft Skin Adhesive, DuPont; mixed 1:1 A:B) and PDMS (RTV-615; mixed 10:1) at a weight ratio of 40:1 (Adhesive:PDMS). The method was based on Chu et. al [33].

For the replaceable substrate layer, a glass slide was plasma-treated for 60 s, spin-coated with the hybrid polymer at 2500 rpm for 1 min and cured at 50°C overnight. This substrate was aligned with the permanently bonded PDMS layers and gently pressed to eliminate trapped air bubbles.

For the replaceable 3D chambers, positive molds were designed in Onshape, processed using Utility Version 6.3.0, and fabricated using a high-resolution 3D printer (285D, CADworks3D). Printed molds were rinsed with isopropanol, air-dried to clear fine features, and post-cured under UV light (405 ± 5 nm). The chamber structure was cast using 10:1 PDMS and cured overnight at 80°C. The surface was subsequently spin-coated with the hybrid polymer (2500 rpm, 1 min) and cured at 50°C overnight.

### Device clean up and reuse

To regenerate the device between trials, ultrapure water and air were sequentially pumped through all microfluidic channels. The PDMS layer was detached from the used substrate, then subjected to a secondary rinse, ultrasonication, and optional autoclaving. The PDMS layer was dried using an air gun and dehydrated in a 60 °C oven for at least 30 minutes. Reassembly was achieved by bonding a fresh substrate to the PDMS layer, securing the device between a metal tray and an acrylic lid, and connecting the control pins.

### Burst test

The bond pressure test was made for 5 different conditions: PDMS device for glass substrate (negative control), spin-coat PDMS with 10:1 or 20:1 ratio, and spin-coat hybrid polymer with 50°C or 80°C curing temperature. These glass substrates were aligned under the permanently bonded PDMS layers and gently pressed to eliminate trapped air bubbles. The control pressure was maintained at 10 psi and the fluid inlet pressure was controlled through pressure-relief vials via PEEK tubing. While all control valves were closed, the fluid inlet flow was controlled through an air regulator to gradually increase the pressure until delamination happens between the fluid layer and the bottom substrate.

### Device functionality test

In order to test the functionality of applying negative pressure, a vacuum source with –2.9 psi (−20kPa) was connected in one outlet. The control pressure was set to 15 psi and fluid pressure was set to 2 psi for chip operation. Before loading food dye, the device was either loaded with Pluronic F-127 0.2% based on the method from Gomez et al [10]; or coated with Teflon AF-2400 0.5% by flowing this solution throughout the fluid channels and put in vacuum chamber for 20 mins to vaporize all extra reagents [35].

### Microfluidic experiments with macrophage

Microfluidic chambers were coated with 250ug/mL fibronectin (R&D Systems) overnight, followed by a wash with fresh medium. Macrophages were loaded into the diffusion chambers at approximately 80% confluency. Cells were allowed to attach and spread for at least 3 hours, with 1–2 media exchanges during this period. Any cells settled in non-chamber channels were removed by flushing with trypsin.

Fluid handling was controlled by a custom MATLAB GUI adapted from Gómez-Sjöberg et al. [10], which managed automated feeding, sinking, and purging cycles. To characterize NF-κB dynamics, cells were stimulated with LPS at concentrations ranging from 0.03 to 30 ng/mL. Stimulation protocols included exponentially increasing doses or random patterns administered at intervals of 30 min, 1 h, 1.5 h, 2 h, or 4 h, controlled by an automated MATLAB scheduler.

### Image acquisition and analysis

Time-lapse fluorescence imaging was performed using a Leica Thunder system with a 20× objective. Images were acquired every 6 minutes using LasX software (Leica). The p65-GFP reporter was imaged using 475 nm excitation (500 ms exposure, 100% LED intensity). Nuclei (DAPI) were imaged using 395 nm excitation (100 ms exposure, 50% intensity).

Post-processing included dark frame subtraction and flat-field correction for all images.

## Acknowledgement

The study is partially supported by the Chan Zuckerberg BioHub Spoke Award and VinUniversity Fellowship from VinUni-Illinois Smart Health Center, VinUniversity.

## Reference

[1] Vyawahare, Saurabh, Andrew D. Griffiths, and Christoph A. Merten. “Miniaturization and parallelization of biological and chemical assays in microfluidic devices.” Chemistry & biology 17.10 (2010): 1052–1065.

[2] Bai, Yang, et al. “Applications of microfluidics in quantitative biology.” Biotechnology Journal 13.5 (2018): 1700170.

[3] Zhong, Qifeng, et al. “Advances of microfluidics in biomedical engineering.” Advanced materials technologies 4.6 (2019): 1800663.

[4] Yang, Yong, et al. “Microfluidics for biomedical analysis.” Small methods 4.4 (2020): 1900451.

[5] Sackmann, Eric K., Anna L. Fulton, and David J. Beebe. “The present and future role of microfluidics in biomedical research.” Nature 507.7491 (2014): 181–189.

[6] Shembekar, Nachiket, et al. “Droplet-based microfluidics in drug discovery, transcriptomics and high-throughput molecular genetics.” Lab on a Chip 16.8 (2016): 1314–1331.

[7] Wong, Ada Hang-Heng, et al. “Drug screening of cancer cell lines and human primary tumors using droplet microfluidics.” Scientific reports 7.1 (2017): 9109.

[8] Song, Helen, Delai L. Chen, and Rustem F. Ismagilov. “Reactions in droplets in microfluidic channels.” Angewandte chemie international edition 45.44 (2006): 7336–7356.

[9] Makgwane, Peter Ramashadi, and Suprakas Sinha Ray. “Synthesis of nanomaterials by continuous-flow microfluidics: a review.” Journal of Nanoscience and Nanotechnology 14.2 (2014): 1338–1363.

[10] Gómez-Sjöberg, Rafael, et al. “Versatile, fully automated, microfluidic cell culture system.” Analytical chemistry 79.22 (2007): 8557–8563.

[11] Kellogg, Ryan A., et al. “High-throughput microfluidic single-cell analysis pipeline for studies of signaling dynamics.” Nature protocols 9.7 (2014): 1713–1726.

[12] Lecault, Véronique, et al. “High-throughput analysis of single hematopoietic stem cell proliferation in microfluidic cell culture arrays.” Nature methods 8.7 (2011): 581–586.

[13] Low, Lucie A., et al. “Organs-on-chips: into the next decade.” Nature Reviews Drug Discovery 20.5 (2021): 345–361.

[14] Bhatia, Sangeeta N., and Donald E. Ingber. “Microfluidic organs-on-chips.” Nature biotechnology 32.8 (2014): 760–772.

[15] Skardal, Aleksander, et al. “Multi-tissue interactions in an integrated three-tissue organ-on-a-chip platform.” Scientific reports 7.1 (2017): 8837.

[16] Guo, Mira T., et al. “Droplet microfluidics for high-throughput biological assays.” Lab on a Chip 12.12 (2012): 2146–2155.

[17] Eyer, Klaus, et al. “A microchamber array for single cell isolation and analysis of intracellular biomolecules.” Lab on a Chip 12.4 (2012): 765–772.

[18] Ma, Zhen, et al. “Self-organizing human cardiac microchambers mediated by geometric confinement.” Nature communications 6.1 (2015): 7413.

[19] Melin, Jessica, and Stephen R. Quake. “Microfluidic large-scale integration: the evolution of design rules for biological automation.” Annu. Rev. Biophys. Biomol. Struct. 36.1 (2007): 213–231.

[20] Gong, Hua, Adam T. Woolley, and Gregory P. Nordin. “High density 3D printed microfluidic valves, pumps, and multiplexers.” Lab on a Chip 16.13 (2016): 2450–2458.

[21] Watson, Craig, et al. “Multiplexed microfluidic chip for cell co-culture.” Analyst 147.23 (2022): 5409–5418.

[22] Mehling, Matthias, and Savaş Tay. “Microfluidic cell culture.” Current opinion in Biotechnology 25 (2014): 95–102.

[23] Jenkins, Gareth, and Colin D. Mansfield, eds. Microfluidic diagnostics: methods and protocols. Totowa, NJ, USA:: Humana Press, 2013.

[24] Payne, Emory M., et al. “High-throughput screening by droplet microfluidics: perspective into key challenges and future prospects.” Lab on a Chip 20.13 (2020): 2247–2262.

[25] Lee, Seok Woo, and Seung S. Lee. “Shrinkage ratio of PDMS and its alignment method for the wafer level process.” Microsystem Technologies 14.2 (2008): 205–208.

[26] Moraes, Christopher, Yu Sun, and Craig A. Simmons. “Solving the shrinkage-induced PDMS alignment registration issue in multilayer soft lithography.” Journal of micromechanics and microengineering 19.6 (2009): 065015.

[27] Madsen, Morten Hannibal, et al. “Accounting for PDMS shrinkage when replicating structures.” Journal of Micromechanics and Microengineering 24.12 (2014): 127002.

[28] Kamei, Ken-ichiro, et al. “3D printing of soft lithography mold for rapid production of polydimethylsiloxane-based microfluidic devices for cell stimulation with concentration gradients.” Biomedical microdevices 17.2 (2015): 36.

[29] Shakeri, Amid, Shadman Khan, and Tohid F. Didar. “Conventional and emerging strategies for the fabrication and functionalization of PDMS-based microfluidic devices.” Lab on a Chip 21.16 (2021): 3053–3075.

[30] Liu, Dong, and Suresh V. Garimella. “Investigation of liquid flow in microchannels.” Journal of Thermophysics and heat transfer 18.1 (2004): 65–72.

[31] Koo, Junemo, and Clement Kleinstreuer. “Liquid flow in microchannels: experimental observations and computational analyses of microfluidics effects.” Journal of Micromechanics and Microengineering 13.5 (2003): 568.

[32] Raj M, Kiran, and Suman Chakraborty. “PDMS microfluidics: A mini review.” Journal of Applied Polymer Science 137.27 (2020): 48958.

[33] Chu, M., et al. “Plasma free reversible and irreversible microfluidic bonding.” Lab on a Chip 17.2 (2017): 267–273.

[34] Mukhopadhyay, Rajendrani. “When PDMS isn’t the best.” (2007): 3248–3253.

[35] Cho, Sung Hwan, Jessica Godin, and Yu-Hwa Lo. “Optofluidic waveguides in Teflon AF-coated PDMS microfluidic channels.” IEEE Photonics Technology Letters 21.15 (2009): 1057–1059.

[36] Son, Minjun, et al. “NF-κB responds to absolute differences in cytokine concentrations.” Science signaling 14.666 (2021): eaaz4382.

[37] Wang, Andrew G., et al. “NF-κB memory coordinates transcriptional responses to dynamic inflammatory stimuli.” Cell reports 40.7 (2022).

[38] Wang, Andrew G., et al. “Macrophage memory emerges from coordinated transcription factor and chromatin dynamics.” Cell Systems 16.2 (2025).

[39] Hoffmann, Alexander, and David Baltimore. “Circuitry of nuclear factor κB signaling.” Immunological reviews 210.1 (2006): 171–186.

[40] Sen, Supriya, et al. “Gene regulatory strategies that decode the duration of NFκB dynamics contribute to LPS-versus TNF-specific gene expression.” Cell systems 10.2 (2020): 169–182.

[41] Lane, Keara, et al. “Escalating threat levels of bacterial infection can be discriminated by distinct MAPK and NF-κB signaling dynamics in single host cells.” Cell Systems 8.3 (2019): 183–196.

[42] Cheng, Quen J., et al. “NF-κB dynamics determine the stimulus specificity of epigenomic reprogramming in macrophages.” Science 372.6548 (2021): 1349–1353.

[43] Kellogg, Ryan A., et al. “Digital signaling decouples activation probability and population heterogeneity.” elife 4 (2015): e08931.

{44] DeFelice, Mialy M., et al. “NF-κB signaling dynamics is controlled by a dose-sensing autoregulatory loop.” Science signaling 12.579 (2019): eaau3568.

[45] Son, Minjun, et al. “Processing stimulus dynamics by the NF-κB network in single cells.” Experimental & Molecular Medicine 55.12 (2023): 2531–2540.

[46] Son, Minjun, et al. “Spatiotemporal NF-κB dynamics encodes the position, amplitude, and duration of local immune inputs.” Science Advances 8.35 (2022): eabn6240.

[47] Fischer, Sarah CL, et al. “Adhesion and Cellular Compatibility of Silicone-Based Skin Adhesives.” Macromolecular Materials and Engineering 302.5 (2017): 1600526.

[48] Boyadzhieva, Silviya, et al. “Thin film composite silicon elastomers for cell culture and skin applications: manufacturing and characterization.” Journal of Visualized Experiments: JoVE 137 (2018): 57573.

